# An ONT sequencing framework for hantavirus genome assembly, variant analysis, and defective viral genome characterization from whole-segment amplicons

**DOI:** 10.64898/2026.10.19.760236

**Authors:** Léna Vandenabeele, Frances M. Twohig, Anne Tuiskunen Bäck, Magnus Sjöblom, Rebecca Lantto, Gabriella Svala, Janne Tynell, Sandra Broddesson, Robert Dyrdak, Wanda Christ, Silke Riesle-Sbarbaro, Joseph B. Prescott, Steven B. Bradfute, Daniel P. Depledge, Benjamin E. Nilsson-Payant

## Abstract

Orthohantaviruses are zoonotic, tri-segmented negative-sense RNA viruses of rodents and other small mammals that can cause severe human disease. Whole genome sequencing remains difficult because of low viral loads, high sequence diversity, and an inherent instability of the viral L segment. Current tiled amplicon and hybrid-capture approaches do not capture the heterogeneity of virus populations, including defective viral genomes (DVGs), which have never been described for hantaviruses. Here, we present an Oxford Nanopore Technologies sequencing framework based on universal reverse transcription primers targeting the conserved genome termini of orthohantaviruses and a single species-specific primer pair per segment, enabling amplification and sequencing of complete S, M, and L segments. We also developed an accompanying bioinformatic pipeline that performs reference-guided assembly of hantavirus genomes, consensus generation, variant calling, and DVG classification using the newly developed ORCHID classification pipeline. We applied this approach to diverse Puumala virus strains from infected cell cultures and bank voles, to representative Old World and New World orthohantaviruses, and to highly pathogenic New World viruses in vivo. In all cases we were able to accurately reconstruct full hantavirus genomes. Notably, we found that hantavirus populations contain highly abundant DVGs and conserved L segment deletions, whereas the S and M segments remained largely intact. Together, this work provides a low-cost, broadly applicable and easily customisable workflow for hantavirus genomics and reveals DVGs as a previously unrecognized feature of hantavirus populations.

**IMPORTANCE:** Hantaviruses cause severe and often fatal human diseases, and their diversity and low viral loads make them hard to sequence, which limits surveillance and outbreak investigation. We describe a simple, universal but customisable, low-cost nanopore sequencing workflow that recovers complete genome segments, including the full-length L segment, from cell culture and animal tissues across the entire *Orthohantavirus* genus. Using this approach, we report the first evidence of defective viral genomes in hantaviruses. These truncated genomes arise mainly from the L segment, which encodes the viral polymerase, and are formed at conserved genomic regions. Defective viral genomes can modulate viral replication, innate immune activation, and persistence in other viruses. Their presence in hantaviruses raises new questions about polymerase function, tolerance in reservoir hosts, and immune-mediated disease in humans. Our framework offers a practical tool for hantavirus genomics and a foundation for investigating how defective genomes shape hantavirus infection.

## INTRODUCTION

Orthohantaviruses (family *Hantaviridae*, class *Bunyaviricetes*) are zoonotic segmented negative-sense RNA viruses that are naturally found in small mammal species, including rodents, shrews, moles and bats (1). Human infections occur primarily through inhalation of aerosolized excreta from infected animal hosts and can cause severe disease, including haemorrhagic fever with renal syndrome (HFRS) and nephropathia epidemica (NE) caused by Old World hantaviruses primarily in Eurasia and hantavirus cardiopulmonary syndrome (HCPS) caused by New World hantaviruses in the Americas (2, 3). Case fatality rates for hantavirus diseases vary significantly depending on the infecting virus species (4): it ranges from <1% for Puumala virus (PUUV), the causative agent for NE, to up to 15% for HFRS-causing Old World hantaviruses such as Hantaan virus (HTNV) in East Asia and Dobrava-Belgrade virus (DOBV) in eastern Europe, and over 30% for HCPS-causing New World hantaviruses such as Andes virus (ANDV) and Sin Nombre virus (SNV).

Hantaviruses have small tripartite negative-sense RNA genomes comprising the small (S), medium (M) and large (L) genome segments, encoding only 4-5 viral proteins (5). The S segment encodes the nucleoprotein (NP) and in some cases a non-structural (NS) protein, the M segment encodes the glycoprotein precursor (GPC) which is co-translationally cleaved into the Gn and Gc glycoproteins, and the L segment encodes the RNA-dependent RNA polymerase (RdRp). The genetic sequence divergence at the amino acid level is significant across the hantavirus family (up to 75%) and even within a species (up to 24%) (6). Low viral RNA copy numbers in clinical and rodent samples, and natural RNA degradation outside of infected hosts makes obtaining complete coverage of hantavirus genomes challenging (7, 8). Current state-of-the-art sequencing approaches include metagenome sequencing, tiled amplicon sequencing and hybrid capture-based sequencing, which all require extensive optimisation and large numbers of probes (9–12). Existing long-read amplicon-based sequencing approaches often include single amplicons for the smaller segments, but to date, there are no established protocols to reliably capture the entire L segment in a single amplicon. In addition, while these methods can provide full genome coverage sequencing and provide consensus sequences of viruses, they do not accurately reflect the genetic makeup of any given virus population. In reality, virus populations are highly heterogeneous and contain large proportions of defective viral genomes (DVGs) consisting of genome truncations, duplications and recombinations (13). However, DVGs have so far never been reported for hantaviruses.

Here, we designed a whole segment amplicon-based long-read nanopore sequencing method and an end-to-end bioinformatical analysis pipeline for hantavirus genome assembly, alignment, consensus sequence generation, variant calling and defective viral genome analysis (Fig.1A). This approach minimises required optimisation steps for tiling and hybridization probes and enables a much more accurate depiction of heterogenous virus populations without sacrificing sensitivity or genome coverage. Using this method, we were able to sequence several diverse PUUV strains from both infected cell cultures and from bank voles – their natural host species. In addition, we validated our method using representative members of the whole *Orthohantavirus* genus and samples obtained from cell cultures and animal tissues. Our sequencing approach provides a cost-effective and rapid end-to-end pipeline to analyse hantavirus genomes from environmental, animal and cellular samples. In addition, we provide the first report of hantavirus DVGs and provide a framework for their analysis.

**FIG 1:**
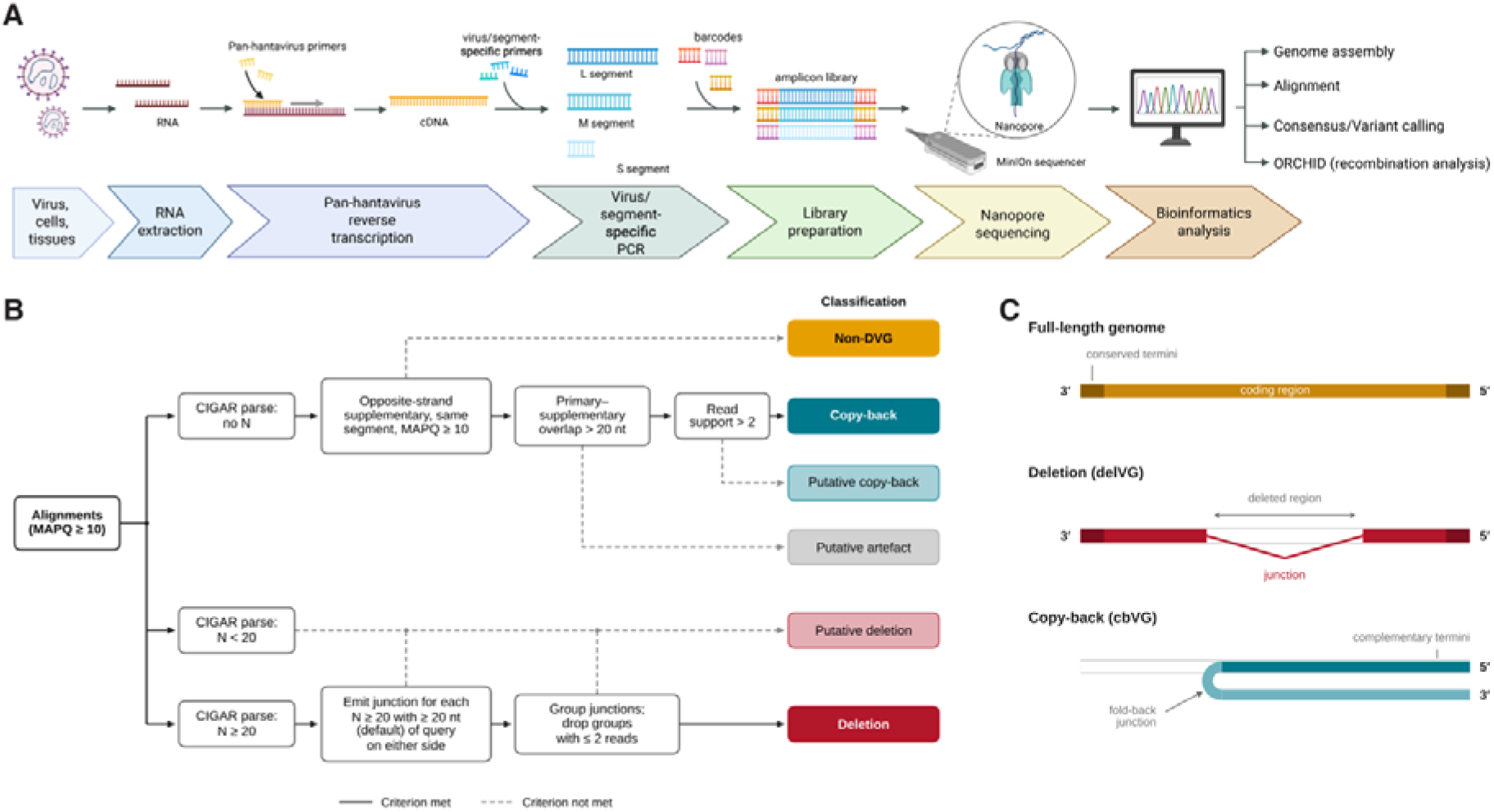
Whole-segment amplicon sequencing workflow and ORCHID classification of defective viral genomes. (A) Schematic of the vgsRT-PCR nanopore sequencing workflow. RNA extracted from virus stocks, infected cells, or tissues is reverse transcribed with pan-hantavirus primers complementary to the conserved genome termini. Each genome segment (L, M, and S) is then separately amplified as a single amplicon with virus- and segment-specific primers, barcoded, pooled into an amplicon library, and sequenced on an Oxford Nanopore Technologies (ONT) MinION device. Reads are used for genome assembly, alignment, consensus and variant calling, and defective viral genome (DVG) analysis with ORCHID (ONT-based Recombination and Copy-back Haplotype Identification for DVGs). Schematic created in BioRender: Nilsson-Payant, B. (2026) https://BioRender.com/0whr19b (B) Decision tree used by ORCHID to classify aligned reads (mapping quality [MAPQ] ≥10). Reads without N operations in their CIGAR string are tested for an opposite-strand supplementary alignment on the same segment. Depending on primary–supplementary overlap and read support, they are classed as copy-back, putative copy-back, or putative artefact. Reads with N operations shorter than 20 nt are classed as putative deletions. For reads with N operations of ≥20 nt, a junction is emitted if ≥20 nt of query aligns on each side. Junctions are grouped across reads, and groups with insufficient read support are dropped. Reads that meet none of the criteria are classed as non-DVG. Solid lines, criterion met; dashed lines, criterion not met. Thresholds are given in Materials and Methods. (C) Schematic of the genome architectures detected. Full-length genomes carry conserved termini flanking the coding region. Deletion DVGs (delVGs) contain an internal deletion, with the junction marking where the polymerase disengaged from and reengaged the template. Copy-back DVGs (cbVGs) arise when the polymerase reengages the nascent strand at a fold-back junction, generating complementary termini.

## RESULTS

### A nanopore sequencing and bioinformatics pipeline

Here, we designed a whole segment amplicon-based long-read nanopore sequencing method. Hantavirus RNA is reverse transcribed using universal hantavirus primers that hybridize with the highly conserved 14 nucleotides at the genome termini. Subsequently, each genome segment is amplified in a virus species-specific manner using a single longer primer pair corresponding to the specific genome termini of a virus genome segment. PCR products are then sequenced using Oxford Nanopore Technologies (ONT) long-read sequencing technology. This is followed by a bioinformatical analysis pipeline that includes hantavirus reference-guided genome assembly, genome alignment, consensus sequence generation, variant calling and DVG analysis (Fig.1A). The DVG analysis is performed using the ‘ONT-based Recombination and Copy-back Haplotype Identification for DVGs’ (ORCHID) pipeline which detects recombination events in virus genomes and classifies these as artefacts, deletion, or copy-back events (Fig. 1B). Deletion DVGs (delVGs) are produced when the RdRp dissociates from the genome template during replication, continues reading the template without copying any of it, and then reassociates and starts copying the template further downstream, thus creating an internal deletion of the copied genome (Fig. 1C). Copy-back DVGs (cbVGs) are produced when the RdRp dissociates from the template strand during replication and reengages the nascent product strand as a new template (Fig. 1C). Together, this workflow represents a highly customisable end-to-end sequencing and analysis framework for hantaviruses.

### Establishment of whole hantavirus segment amplification

While amplification of the S and M segments in single amplicons has been achieved before, capturing the entire L segment in one single amplicon has so far proven difficult (10). We utilized the extremely high sequence conservation in the terminal 14 nucleotides of the 5’ and 3’ ends of all three genome segments across the entire *Orthohantavirus* genus to design universal hantavirus reverse transcription primers for both genomic negative-sense viral RNA (vRNA) and the antigenomic positive-sense complementary RNA (cRNA) (Fig. 2A). Primers for amplification of individual segments also contained these highly conserved termini but stretched further into the segments to achieve high specificity for virus species and segments.

**FIG 2:**
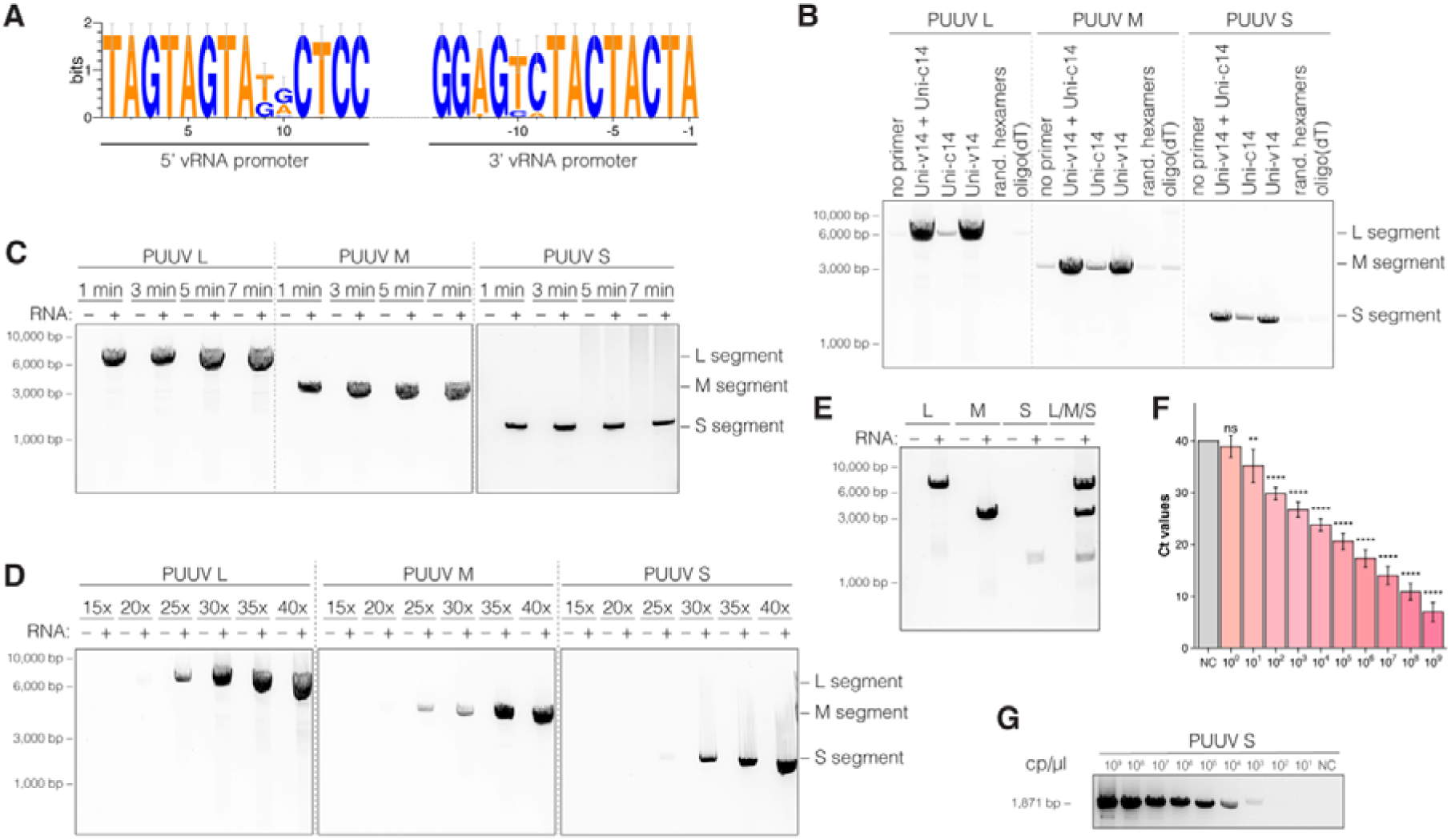
Establishment and optimization of whole-segment vgsRT-PCR for Puumala virus (PUUV). (A) Sequence logo of the 14 terminal nucleotides at the 5′ and 3′ ends of the genomic vRNA segments of 33 diverse *Orthohantavirus* genomes. Letter height indicates information content (bits); positions are numbered from the respective genome end. (B-E) RNA from a purified PUUV Kazan-E6 stock was reverse transcribed followed by segment-specific PCR (vgs-RT-PCR) and resolved by agarose gel electrophoresis. (B) Different RT primers, (C) PCR elongation times, (D) PCR cycle numbers and (E) primer multiplexing were assessed. (F) RT-qPCR cycle threshold (Ct) values for a 10-fold dilution series of *in vitro*-transcribed PUUV S segment vRNA. NC, negative control. Data represent the mean ± the standard deviation of 4 replicates. Statistical significance was determined by one-way ANOVA followed by Dunnett’s correction for multiple comparisons: **** (p < 0.0001), *** (p < 0.001), ** (p < 0.01), * (p < 0.05), ns (p > 0.05) (G) vgsRT-PCR amplification of the PUUV S segment from the same dilution series.

We established and optimised our virus species- and segment-specific viral genome segment RT-PCR (vgsRT-PCR) method on RNA extracted from a cell-free purified PUUV stock. First, we compared different RT primers and found that the Uni-c14 primer, random hexamers and oligo(dT) primers performed poorly and resulted in low or no detectable amplification of PUUV segments (Fig. 2B). This was an expected result, since hantaviruses generally are not thought to contain polyadenylation signals and since the virus stock primarily should be composed of vRNA (14, 15). However, use of the vRNA-specific Uni-v14 primer alone or in combination with the cRNA-specific Uni-c14 primer yielded strong full-length amplification of all three PUUV segments. Henceforth, the combination of vRNA- and cRNA-specific RT primers was used. Next, different PCR elongation times and cycle numbers were compared. We observed that for the longer L and M segments, increasing elongation time seemed to slightly increase full-length segment amplification, while no major difference was observed for the S segment (Fig. 2C). Based on these data we selected an elongation time of 3 min as the best balance between sensitivity and overall reaction time. Increasing PCR cycle numbers steadily increased PCR product accumulation in all three segments, with robust product accumulation starting at 30 cycles (Fig 2D). As a compromise between sensitivity, PCR bias and time optimisation, we chose 30 cycles as our default option, with the possibility of increasing cycles to 40 for samples that require higher sensitivity. As a proof-of-concept we also tested whether all segment-specific primers can be combined in a single reaction and confirmed that all PUUV segments were efficiently and equally amplified compared to the single primer pair reactions (Fig. 2E). Sensitivity is important when designing PCR protocols for hantaviruses due to their inherently low RNA copy numbers in infected patients. Using *in vitro* transcribed PUUV S segment vRNA as a standard, we were able to accurately detect up to 10^1^ RNA copies by RT-qPCR (Fig. 2F) and up to 10^3^ RNA copies by vgsRT-PCR (Fig. 2G).

Overall, we established a reproducible and sensitive vgsRT-PCR method for the optimal amplification of whole PUUV segments with a single primer pair per segment.

### Nanopore sequencing of diverse PUUV strains from cell culture

Having established that we can robustly amplify full-length PUUV genomes, we next attempted to sequence and characterise these amplicons from more complex samples by ONT long-read sequencing. To that end, we infected human lung epithelial cells with several different PUUV strains and extracted RNA from infected cells. First, we confirmed and compared the presence of viral RNA by RT-qPCR and found robust – but varying – levels of viral RNA in all samples (Fig.3A). Analysis of vgsRT-PCR products by gel electrophoresis demonstrated clean amplification of the full M and S segments for all PUUV strains (Fig. 3B). However, for the L segment we predominantly observed PCR products of varying shorter lengths, suggesting production of internally truncated DVGs that maintained the conserved segment termini. This was especially pronounced for the PUUV strain BE/Cg/Villers-la-Tour/1/2009 (‘Villers-la-Tour’), BE/Cg/Seloignes/1/2009 (‘Seloignes’), BE/Cg/Beaumont/1/2012 (‘Beaumont’), and BE/Cg/Turnhout/1/1985 (‘Turnhout’), while RUS/Cg/Kazan/E6/1982 (‘Kazan-E6’) exhibited relatively clean full-length products. Interestingly, while these putative DVGs exhibited strain-specific profiles, the overall size distribution appears similar across all virus strains.

**FIG 3:**
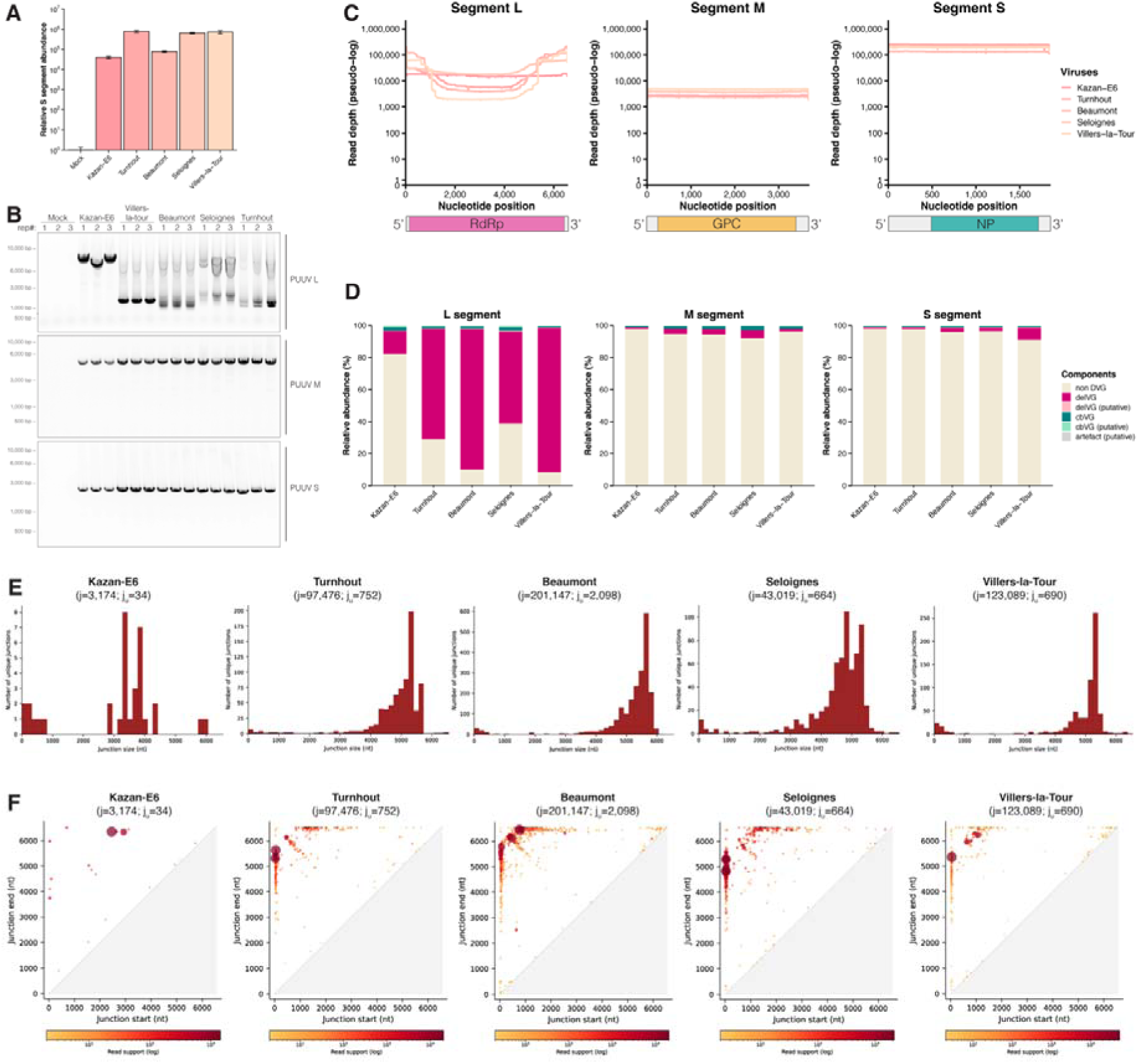
Sequencing of diverse PUUV strains from infected cell culture reveals abundant L segment deletion DVGs. A549 cells were infected with the indicated PUUV strains (MOI = 0.1; 72 hpi) or mock infected. (A) RT-qPCR quantification of PUUV S segment RNA, calculated by the ΔΔCt method relative to *ACTB* and normalized to uninfected controls. Data represent the mean ± the standard deviation of 3 replicates. (B) vgsRT-PCR of the L, M, and S segments from three independent infections, resolved by agarose gel electrophoresis. (C) Nanopore read depth (pseudo-log scale) across the L, M, and S segments. Each line represents one strain. Schematics below show the coding and untranslated regions. (D) ORCHID read classification for each segment, shown as relative abundance (%). (E) Size distribution of unique L segment delVG junctions (length of the deleted region in nt). j, total number of junction-supporting reads; j_u_, number of unique junctions. (F) Junction start and end coordinates of L segment delVGs. Point size and color indicate read support (log scale).

Nanopore sequencing of these amplicons demonstrated high levels of sequencing depth across the entire genome (Table 1 and Fig. 3C). The genome coverage was even across the entire genome for the M and S segments for all PUUV strains, confirming our previous vgsRT-PCR data. Accordingly, we observed a striking reduction in genome coverage in the central regions of the L segment for all strains except Kazan-E6, which also did not generate any major truncated PCR products. We used the ORCHID pipeline to classify sequenced viral reads and confirmed our vgsRT-PCR and genome coverage data: while the M and S segments only contained small fractions of putative and high-confidence DVGs, the vast majority of L segment reads were classified as delVGs (Table 1 and Fig. 3D). Only Kazan-E6 exhibited a lower – yet still significant – fraction of DVGs, consistent with our previous findings. As the M and S segments did not exhibit any meaningful levels of DVGs, and cbVGs could not be detected at significant levels in any segment, we characterized only L segment delVGs further. Analysis of L segment delVG junction sizes revealed a now familiar pattern: while Kazan-E6 only showed few unique junctions that were concentrated between 3000 and 4000 nt in size, all the remaining strains almost exclusively had very large (approximately 4000 to 6000 nt in size) deletions (Fig. 3E). This helps explain why we previously observed a relatively uniform pattern of truncated PCR products (Fig. 3B). Investigation of the exact junction sites – *i.e.* at what nucleotide position of the genome the RdRp disengages from the template (junction start) and then subsequently reengages again (junction end) – helps shed further light onto the nature of these DVGs and reveals highly specific regions in the PUUV L segment that are associated with delVG formation (Fig. 3F). For the Turnhout, Beaumont, Seloignes and Villers-la-Tour strains we observed highly similar patterns: the majority of deletions start within the first 100 nt of the 5’ end of the L segment and span almost the entire segment and end within a more variable (approximately 1500 nt) window at the 3’ end of the segment (Fig.3F).

**Table 1.**
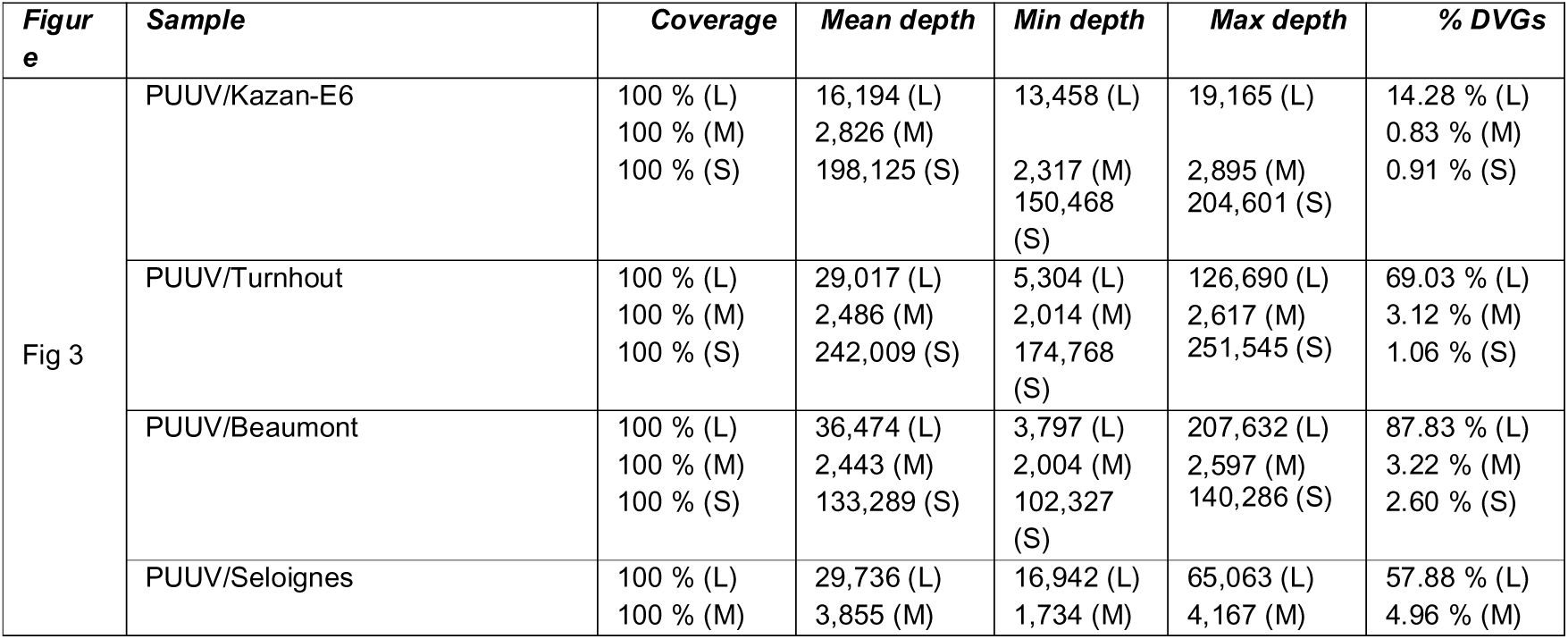
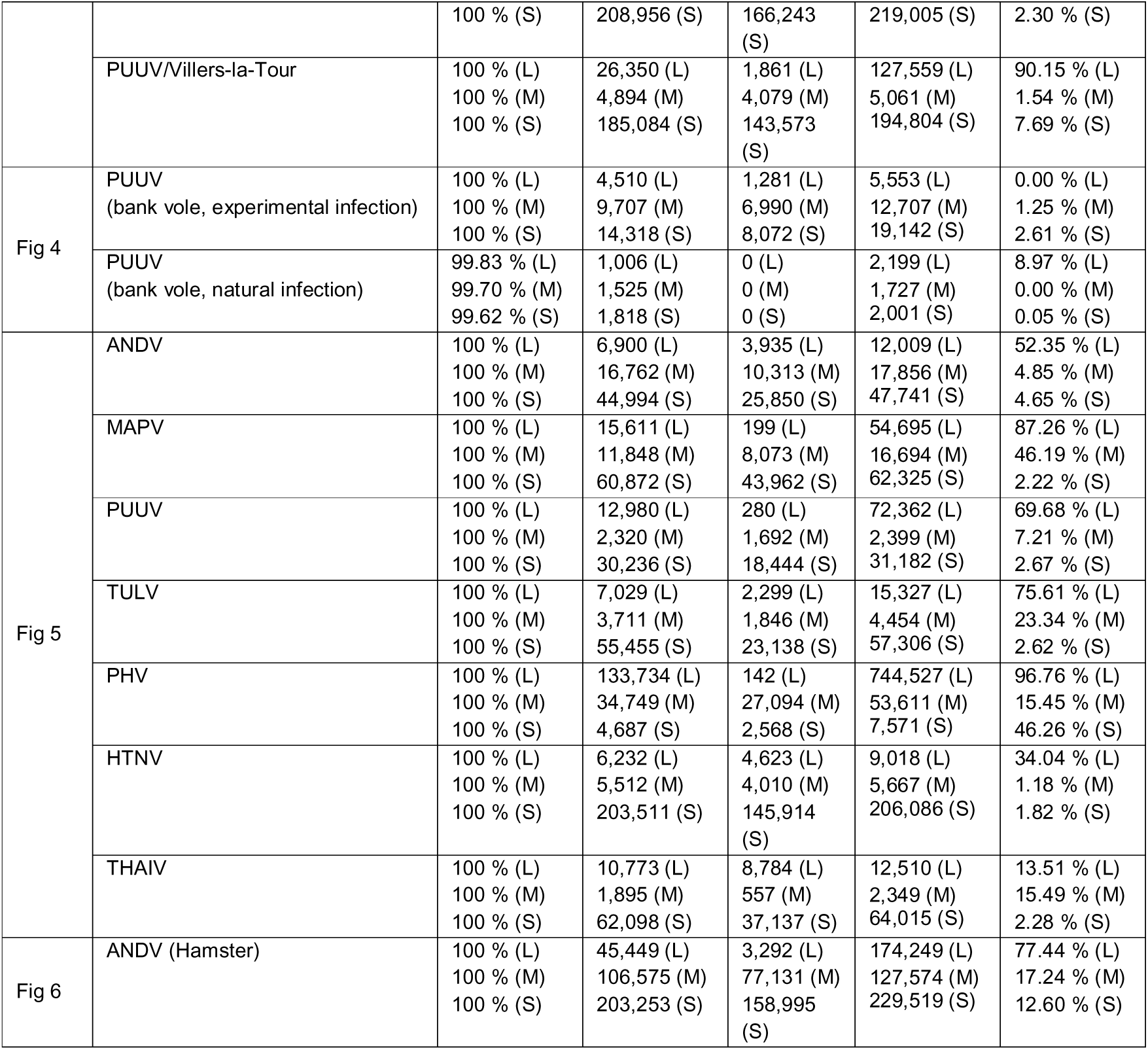
Summary sequencing statistics.

Together, these data demonstrated that our whole segment amplicon sequencing method is able to robustly sequence and resolve entire PUUV genomes. In addition, our analysis framework allows us to confidently call and characterise DVGs from nanopore sequencing data, identifying high abundance delVGs in most PUUV L segments.

### Sequencing of PUUV genomes from bank vole tissues

Having established that we are able to robustly recover full PUUV genomes from cell culture infections, we next attempted to validate this method in highly complex *in vivo* samples. To that end we extracted total RNA from lung tissues obtained either from bank voles experimentally infected with a fully bank vole adapted PUUV strain (16), or from a wild Swedish PUUV-positive bank vole (Fig. 4A). PUUV RNA levels in these samples were low but detectable by RT-qPCR (Fig. 4B). Nanopore sequencing of both wild and experimental infections showed strong genome coverage across all segments (Table 1 and Fig. 4C). Interestingly, no loss of genome coverage could be seen in any segment, including the L segment, with the exception of a putative low frequency deletion in the L segment of the PUUV sequenced from the wild bank vole (Fig. 4C). These observations were fully validated after read classification: almost all sequenced reads could be mapped to the full viral genome, with the most significant – yet small – aberration being L segment delVGs obtained from the wild bank vole isolate (Fig. 4D). Analysis of the junction coordinates for this delVG showed that the start and end points of all delVGs clustered in the same regions close to the 5’ and 3’ genome ends (Fig. 4E).

**FIG 4:**
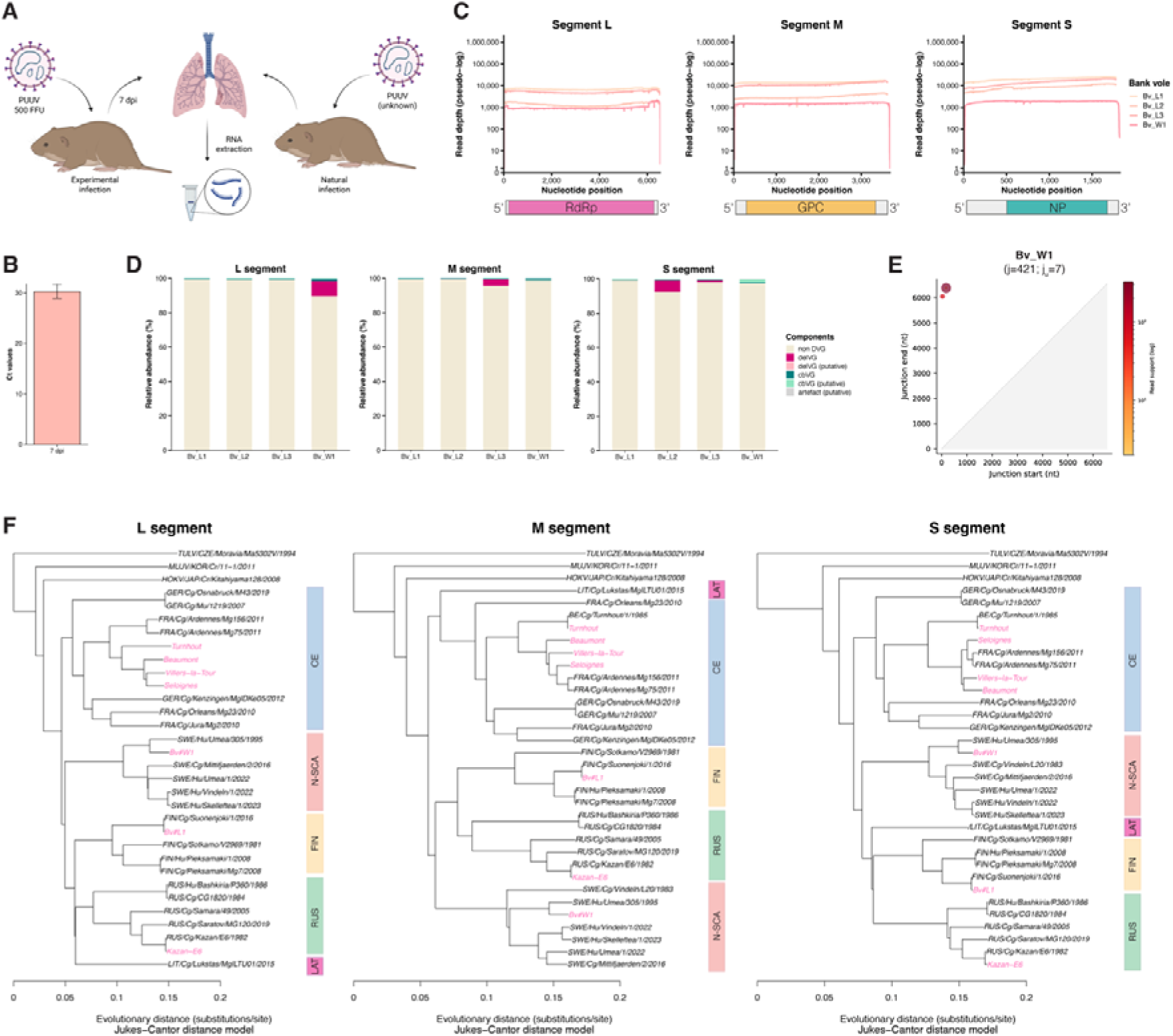
Recovery of PUUV genomes from bank voles. (A) Schematic of bank vole sample collections. Schematic created in BioRender: Nilsson-Payant, B. (2027) https://BioRender.com/rephuh9 (B) RT-qPCR Ct values for PUUV S segment RNA from lungs of bank voles experimentally infected with PUUV strain Suonenjoki and sampled at 7 days postinfection (dpi). Data represent the mean ± the standard deviation from 3 different animals. (C) Nanopore read depth (pseudo-log scale) across the L, M, and S segments for three experimentally infected bank voles (Bv_L1–L3) and one wild-caught PUUV-positive bank vole (Bv_W1). (D) ORCHID read classification for each segment and animal. (E) Junction start and end coordinates of the L segment delVGs detected in Bv_W1. Point size and color indicate read support (log scale); j, total number of junction-supporting reads; j_u_, number of unique junctions. (F) Neighbor-joining phylogenies of L, M, and S segment consensus sequences. Sequences generated in this study are shown in pink, and GenBank reference sequences in black. Tula virus (TULV), Muju virus (MUJV), and Hokkaido virus (HOKV) are included for reference. Colored bars indicate PUUV lineages: CE, Central European; N-SCA, North Scandinavian; FIN, Finnish; RUS, Russian; LAT, Latvian. Segments were aligned with MUSCLE, and trees were built from Jukes–Cantor distances and midpoint-rooted. Scale bar, substitutions per site.

Phylogenetic analysis of the consensus sequences generated here was used as a test to check whether we are able to correctly assemble these PUUV genomes (Fig. 4F). The Kazan-E6 and Turnhout strains clustered together with their known sequences, and the remaining Belgian strains (Beaumont, Seloignes and Villers-la-Tour) all clustered together with the other Belgian and French PUUV strains within the Central European lineage. The Swedish wild bank vole isolate was as expected most closely related to other North-Scandinavian lineage PUUV strains and the experimentally infected bank vole sequences clustered together with the FIN/Cg/Suonenjoki/1/2016 strain used for infections.

Here, we validated our vgsRT-PCR nanopore sequencing pipeline on biologically complex samples from both wild and experimentally infected bank voles and were able to recover high genome coverage, allowing us to confidently predict evolutionary relationships between sequenced PUUV isolates.

### Nanopore sequencing validation of diverse members of the hantavirus family

After having demonstrated that we efficiently can sequence PUUV genomes, we next sought to apply this pipeline to representative members spanning the entire *Orthohantavirus* genus. Since the universal hantavirus RT primers were designed against the entire genus, we subsequently designed virus- and segment-specific primers for the amplification of virus segments. Human A549 cells were infected with a diverse selection of pathogenic and non-pathogenic New World and Old World hantaviruses, including hantaviruses associated with the *Sigmodontinae (*New World mice and rats; ANDV and MAPV), *Arvicolinae* (voles and lemmings; PUUV, TULV and PHV) and *Murinae* (Old World mice and rats; HTNV and THAIV) rodent subfamilies. RNA from infected cells were subsequently analysed by vgsRT-PCR (Fig. 5A) and RT-qPCR (Fig. 5B). Viral RNA was successfully detected in all samples (Fig. 5B). Similar to our results with PUUV, we observed segment-specific differences in DVG production in this panel of hantaviruses, with most DVGs being produced from the L segment (Fig. 5A). Sequencing of hantavirus amplicons confirmed these data (Table 1 and Fig. 5C). L segments consisted mostly of delVG, with only a small proportion of full-length L segments. The M and S segments were mostly constituted of full-length genomes. Two notable exceptions for this were Prospect Hill virus (PHV), which exhibited exceptionally high delVG content in the L (96.76%) and S (46.26%) segments, and Maporal virus (MAPV), which exhibited high delVG content in the L (87.26%) and M (46.19%) segments. Thailand virus (THAIV) - similar to Kazan-E6 previously – had a much lower proportion of DVGs. These findings were reflected in genome coverage across the three virus segments: While genome coverage was high and equal for the M and S segments for all viruses – with the exception of small decreases in coverage in M and S segment coverage for the aforementioned MAPV and PHV, respectively – clear loss of read coverage was seen in the L segment of all viruses with the exception of THAIV, matching our previous findings (Fig. 5D). The L-derived delVGs of all viruses, including the *Murinae*-associated HTNV and THAIV to a lesser extent, exhibited a similar and familiar pattern of junction start and end points within two major clusters (Fig. 5E): one cluster where the junctions start almost immediately following the 5’ end of the L segment and a second cluster where junctions do not start until approximately one third into the L segment. The rejoin point always tends to be towards the far 3’ end of the segment.

**FIG 5:**
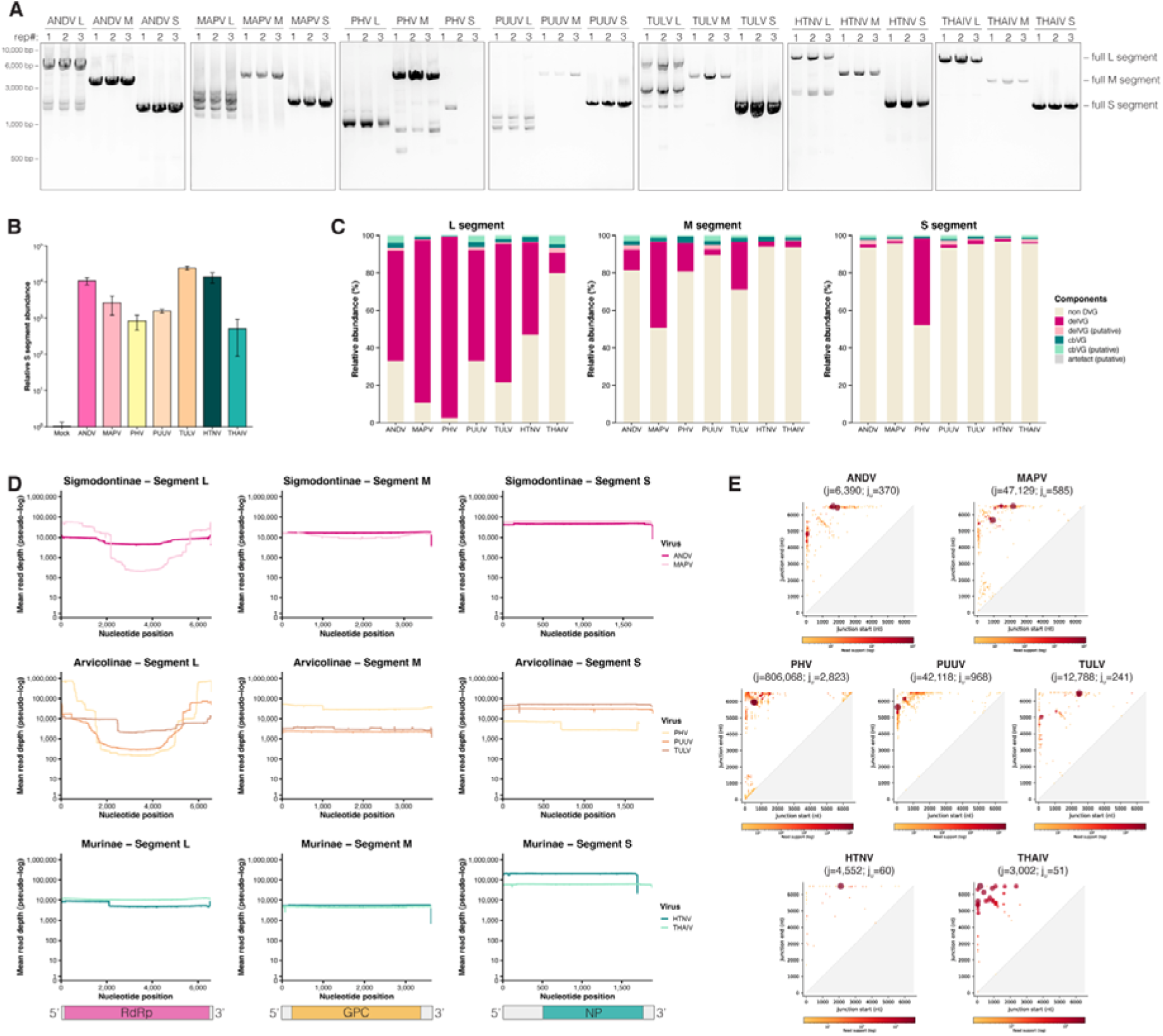
Whole-segment sequencing of representative members of the *Orthohantavirus* genus from infected cells. A549 cells were infected with ANDV, MAPV, PHV, PUUV (strain Turnhout), TULV, HTNV, or THAIV (MOI = 0.1; 96 hpi). (A) vgsRT-PCR of the L, M, and S segments from three independent infections per virus, resolved by agarose gel electrophoresis. (B) RT-qPCR quantification of S segment RNA relative to *ACTB* and normalized to uninfected controls. Data represent the mean ± the standard deviation from 3 independent replicates. (C) ORCHID read classification for each segment and virus. (D) Mean nanopore read depth (pseudo-log scale) across each segment, grouped by the rodent subfamily of the natural reservoir: Sigmodontinae (ANDV, MAPV), Arvicolinae (PHV, PUUV, TULV), and Murinae (HTNV, THAIV). (E) Junction start and end coordinates of L segment delVGs for each virus. Point size and color indicate read support (log scale); j, total number of junction-supporting reads; j_u_, number of unique junctions.

These results confirm that our approach is applicable across the whole *Orthohantavirus* genus and can be used to sequence and analyse diverse hantavirus genomes derived from cell culture.

### Nanopore sequencing of highly pathogenic New World hantaviruses in a relevant *in vivo* human disease model

Syrian hamsters are commonly used as a disease model to mimic HCPS-like disease in humans and challenge with the Chile-9717869 strain of ANDV is uniformly lethal in these animals (17). In order to validate our sequencing method in complex biological samples mimicking human disease, we challenged Syrian hamsters with a lethal dose of ANDV and extracted RNA from lungs of acutely infected animals (Fig. 6A). First, we validated successful infections by RT-qPCR and observed high levels of viral RNA in hamster lungs (Fig. 6B). Analysis by vgsRT-PCR confirmed the clear pattern observed earlier, with efficient and clean detection of the S and M segments, and only weak amplification of full-length L segments and strong accumulation of truncated L-derived DVGs (Fig. 6C). This was confirmed both by observation of genome coverage (Fig. 6D) and by read classification (Fig. 6E and Table 1), showing again significant loss of coverage in the L segment and high proportions of L-derived reads containing deletions. Interestingly, a notable increase in the relative abundance of copy-back and putative copy-back events – although still negligible compared to deletion events - was detected compared to our data from cell culture infections. A closer analysis of the junctions of the DVGs observed in these animals revealed an interesting pattern (Fig. 6F-I). Almost all junctions in delVGs spanned the entire respective genome segment resulting in large junction sizes and small DVG products (Fig. 6F) with junction start and end coordinates clustering close to the 5’ and 3’ termini, respectively (Fig. 6H). However, junctions formed by copy-back events appear very small (Fig. 6G), which can be explained by the fact that the break and rejoin points of the junction are almost exclusively clustered in close proximity, but on opposing orientations of the genome, to each other, resulting in small junction sizes and small DVG products (Fig. 6I). In summary, these data demonstrate that we can also apply our sequencing pipeline to complex human disease models of hantaviruses and efficiently recapture heterogenous hantavirus genome features, including L segment delVGs.

**FIG 6:**
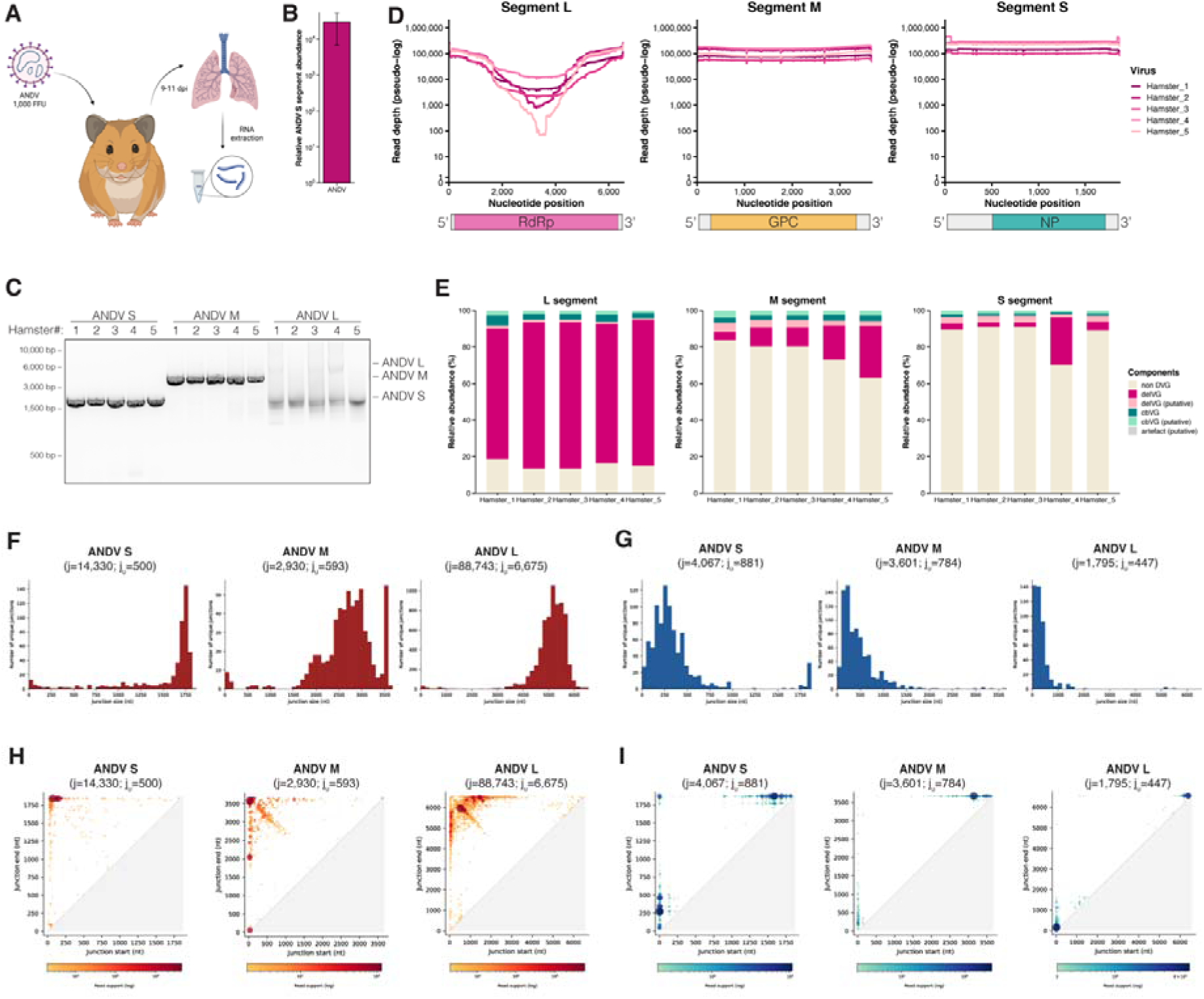
Nanopore sequencing of highly pathogenic New World hantaviruses in an *in vivo* human disease model. (A) Hamsters were intranasally inoculated with 1,000 FFU of ANDV strain Chile-9717869, and lungs were collected at 9–11 dpi for RNA extraction. Schematic was created in BioRender: Nilsson-Payant, B. (2027) https://BioRender.com/ayhnw5x (B) RT-qPCR quantification of ANDV S segment RNA in lung tissue, relative to hamster *ACTB* and normalized to uninfected control animals. Data represent the mean ± the standard deviation of *n* = 5 animals. (C) vgsRT-PCR of the S, M, and L segments from individual hamsters (1–5), resolved by agarose gel electrophoresis. (D) Nanopore read depth (pseudo-log scale) across each segment. Each line represents one hamster. (E) ORCHID read classification for each segment and animal. (F and G) Size distributions of unique delVG (F) and cbVG (G) junctions for each segment (junction size is the length of the deleted region for delVGs and of the shared reference region for cbVGs). (H and I) Junction start and end coordinates for delVGs (H) and cbVGs (I). Point size and color indicate read support (log scale). For cbVGs, junction start and end define the boundaries of the shared reference region. j, total number of junction-supporting reads; j_u_, number of unique junctions.

## DISCUSSION

Here we present an end-to-end workflow for hantavirus genome sequencing and defective viral genome (DVG) analysis, based on whole-segment amplification and nanopore long-read sequencing. By targeting the conserved genome termini of the *Orthohantavirus* genus with universal hantavirus RT primers and amplifying each segment with a single species-specific primer pair, we recovered complete S, M and L segments in all sequenced samples in single amplicons. This has previously been a persistent obstacle for the amplification of full-length L segment in existing protocols (10, 12). The method presented here worked across samples of very different complexity and origin: from *in vitro* transcribed RNA and purified virus stocks, to infected cell cultures, lung tissues from rodents from highly diverse Old World and New World hantaviruses. Furthermore, it was also able to generate phylogenetically coherent consensus sequences, demonstrating that the reference-guided genome assembly retained an unbiasedness within virus species.

Compared to tiled amplicon and hybrid capture approaches, this method needs to tiling scheme or large probe sets and can be adapted to new species by redesigning one primer pair per segment. Its sensitivity of 10^3^ copies based on a gel-based signal is most likely a significant underestimation of the nanopore sequencing sensitivity. Nevertheless, this is highly encouraging for low-titre clinical samples, with previous studies estimating that hantavirus RNA copy numbers can range from 10^3^ to over 10^6^ copies per ml of plasma in NE and HCPS patients, well within range of the sensitivity demonstrated here (7, 8).

An interesting finding was that hantavirus populations contain highly abundant and diverse DVGs, which to our knowledge has not been reported previously. Because the amplicon-based approach captures all individual molecules that contain the conserved genome termini rather than a certain consensus sequence, DVGs were directly visible and measurable in gel electrophoresis, genome coverage profiles and read-level classifications. In nearly all viruses tested, the L segment was the dominant source of DVGs – almost exclusively delVGs – while the M and S segments were largely intact. Copy-back or snap-back DVGs, which are prominent in many non-segmented negative-sense RNA viruses (18–20), were relatively rare, though slightly enriched in ANDV-infected hamster lungs. This pattern is reminiscent of influenza A virus, another segmented negative-sense RNA virus, where internal deletions arise preferentially in the longest segments (PB2, PB1, PA, HA) (21). The size of the L segment might therefore make it particularly prone for deletion events. This could be explained by one of our previous observations: the occurrence of deletions is significantly increased when the supply of available NP becomes scarce (22). This has an immediate impact on the L segment, which is approximately 1.8-fold and 3.6-fold longer than the M and S segments, respectively. However, segment length is unlikely to be the only explanation for the abundance of L segment delVGs, since the break and rejoin coordinates of identified deletions exhibited strong enrichment for particular genomic regions. Across virus strains and species, L-derived delVG junctions fell into discrete clusters: breakpoints began very close to the 5′ end (or, in some viruses, about one-third into the segment) and rejoined near the 3′ terminus, producing large deletions and short delVG products that retain both conserved termini.

DVG abundance differed strongly between different virus strains and species and experimental systems. These differences may reflect several factors, including passage history of the viruses and multiplicity of infection, which have long been known to affect DVG accumulation in viruses (20). However, it is noteworthy that PUUV in its natural reservoir species produced almost no DVGs, whereas ANDV generated high levels of DVGs, including ultra-short delVGs and cbVGs which have lost almost the entire viral segment with the exception of the genome termini, in a human disease animal model. Whether DVGs contribute to, or merely accompany, virus tolerance in reservoir species on the one hand and human disease on the other hand remains open. DVGs can interfere with virus replication, trigger innate immune sensing and promote virus persistence (13, 23, 24), and it will be important to test whether hantavirus DVGs have such activities, particularly given the central role of immune-mediated pathology in HFRS and HCPS (2, 3).

Several limitations should be considered. First, our method requires intact genome termini for amplification, so DVGs lacking one or both termini, or highly fragmented genomes, will not be detected. We argue that genomes lacking the conserved promoter sequences become biologically negligible as they cannot be recognised by the viral RdRp and further amplified. Second, the consensus sequence at the extreme termini is derived from the primers used in this study. Third, RT-PCR generally favours shorter templates, so delVG proportions probably overestimate their true abundance relative to full-length genomes, and the values here should be compared within this workflow rather than taken as absolute frequencies. However, our RT-PCR optimisations demonstrate that under these described reaction conditions any size bias during PCR is negligible (Fig. 2E). Third, PCR and RT template switching can create chimeric products that mimic recombination, which the ORCHID classification can only partly separate. But the reproducibility of junction clusters across independent samples and virus species argues against this as the main source.

In summary, this workflow provides a practical, low-cost approach for generating full-length hantavirus genomes from whole-segment amplicons and for characterising within-sample diversity, including DVGs, from cell culture, animal and clinically relevant samples. The discovery that L-segment delVGs are a conserved feature of hantavirus populations raises new questions about polymerase function, virus–host interactions, persistence and immunopathology. The pipeline we describe here should allow these questions to be addressed in clinically and ecologically relevant samples.

## MATERIALS AND METHODS

### Cell culture

Human alveolar basal epithelial carcinoma cells (A549; ATCC, CCL-185) and African green monkey kidney epithelial cells (Vero E6; ATCC, CRL-1586) were commercially obtained. Bank vole kidney epithelial cell (Mygla.REC.B) were a kind gift by Tomas Strandin (University of Helsinki, Finland) and have previously been described (16). Cells were maintained in Dulbecco’s Modified Eagle Medium (DMEM) supplemented with 10% foetal bovine serum (FBS), L-Glutamine, MEM Non-Essential Amino Acids and 100 U/ml penicillin and 100 μg/ml streptomycin and cultured at 37°C and 5% CO_2_.

### Viruses

Andes virus (ANDV; *Orthohantavirus andesense*) strain Chile-9717869 was a kind gift by Piet Maes (Université Libre de Bruxelles, Belgium). Andes/Chile-9717869 virus was originally isolated from the kidneys and lungs of an infected wild *Oligoryzomys longicaudatus* (long-tailed pygmy rice rat) trapped during an outbreak of HCPS in Chile in 1997 and passaged in Vero E6 cells (25, 26). Maporal virus (MAPV; *Orthohantavirus maporalense*) strain HV9021050 was a kind gift by Piet Maes (Université Libre de Bruxelles, Belgium). Maporal/HV9021050 virus was originally isolated from the spleens of an infected wild *Oligoryzomys fulvescens* (fulvous pygmy rice rat) trapped in Hato Maporal near Caño Delgadito in Venezuela in 1997 and subsequently passaged in Vero E6 cells (27). Hantaan virus (HTNV; *Orthohantavirus hantanense*) strain 76-118 was acquired from the European Virus Archive Global (EVAg) repository (Ref-SKU: 008V-EVA1471) through its partner institution BMC-SAS (Bratislava, Slovakia). Hantaan/76-118 virus was originally isolated from the lungs of an infected wild *Apodemus agrarius* (striped field mouse) in Songnaeri, North Korea, in 1976 and subsequently passaged in A549 cells (28, 29). Thailand virus (THAIV; *Orthohantavirus thailandense*) strain Thai749 was a kind gift by Piet Maes (Université Libre de Bruxelles, Belgium). Thailand/Thai749 virus was originally isolated from the lungs of infected wild *Bandicota indica* (greater bandicoot rat) in Kanchanaburi, Thailand, in 1985 and subsequently propagated in Wistar rats (30). Tula virus (TULV; *Orthohantavirus tulaense*) strain Moravia/Ma5302V/94 was a kind gift by Rainer Ulrich (Friedrich-Loeffler-Institut, Germany). Tula/Moravia/Ma5302V/94 virus was originally isolated from the lungs of infected *Microtus arvalis* (common vole) trapped in Tvrdonice, South Moravia (Czech Republic) in 1994 and subsequently passaged once in a hantavirus-free laboratory *Microtus arvalis* animal and then in Vero E6 cells (31, 32). Prospect Hill virus (PHV; *Orthohantavirus prospectense*) strain PH-1 was acquired from the European Virus Archive Global (EVAg) repository (Ref-SKU: 008V-03727) through its partner institution BMC-SAS (Bratislava, Slovakia). Prospect Hill/PH-1 virus was originally isolated from the lungs of an infected *Microtus pennsylvanicus* (meadow vole) trapped on Prospect Hill in Frederick, Maryland, USA in 1982 and subsequently passaged in Vero E6 cells (33, 34).

Puumala virus (PUUV; *Orthohantavirus puumalaense*) strain RUS/Cg/Kazan/E6/1982 (also known as Kazan-E6) was a kind gift by Jonas Klingström (Linköping University, Sweden). Puumala/RUS/Cg/Kazan/6Cg/1982 (also known as Kazan-wt) virus was originally isolated from the lungs of an infected *Clethrionomys glareolus* (bank vole) trapped in Kazan, Russia, in 1982 and subsequently passaged 21 times in laboratory *C. glareolus* animals (35), before being adapted to Vero E6 cells through at least 24 passages (36, 37). PUUV strain FIN/Cg/Suonenjoki/1/2016 was a kind gift by Tomas Strandin (University of Helsinki, Finland). PUUV/FIN/Cg/Suonenjoki/1/2016 was isolated from the lungs of an infected *C. glareolus* trapped in Suonenjoki, Finland, in 2016 and subsequently passaged in Mygla.REC.B bank vole kidney epithelial cells (16). PUUV strains BE/Cg/Turnhout/1/1985 (previously known as CG13891), BE/Cg/Beaumont/1/2012, BE/Cg/Seloignes/1/2009, and BE/Cg/Villers-la-Tour/1/2009 were a kind gift by Piet Maes (Université Libre de Bruxelles, Belgium). Puumala/BE/Cg/Turnhout/1/1985 virus was isolated from the lungs of an infected *C. glareolus* trapped in Turnhout, Belgium in 1985 and subsequently passaged in Vero E6 cells (38). Puumala/BE/Cg/Beaumont/1/2012 virus was isolated from lungs and kidneys of an infected *C. glareolus* trapped in Beaumont, Belgium in 2012 and subsequently passaged in Vero E6 cells (39). Puumala/BE/Cg/Seloignes/1/2009 virus and Puumala/BE/Cg/Villers-la-Tour virus were isolated from the lungs of infected *C. glareolus* trapped in Seloignes and Villers-la-Tour, Belgium, respectively, in 2009 and subsequently passaged in Vero E6 cells.

All viruses – with the exception of Puumala/FIN/Cg/Suonenjoki/1/2016 virus – were propagated at 37°C in Vero E6 cells in DMEM supplemented with 2% FBS, L-Glutamine, MEM Non-Essential Amino Acids, 100 U/ml penicillin and 100 μg/ml streptomycin as previously described (40). Virus-containing cell culture supernatants were repeatedly collected between 5 to 14 days post-infection (dpi) and replaced with fresh infection media. Pooled viral stocks were purified and concentrated using Amicon Ultra-15 centrifugal filter units (100-kDa molecular weight cut-off). Infectious virus titres were determined by focus-forming assays as described below. Puumala/FIN/Cg/Suonenjoki/1/2016 virus was propagated as previously described in Mygla.REC.B bank vole kidney epithelial cells (16). All cell culture infection were performed in DMEM supplemented with 2% FBS, L-Glutamine, MEM Non-Essential Amino Acids, 100 U/ml penicillin and 100 μg/ml streptomycin.

All cell culture work involving infectious ANDV, MAPV and HTNV was performed in the biosafety level 3 (BSL-3) facility of the Hannover Medical School (MHH) in accordance with its institutional biosafety requirements.

### Focus-forming assay

Infectious viral titres were determined by focus-forming assays as previously described in Vero E6 cells (41). Vero E6 cells were infected with sequential 10-fold dilutions of virus and overlayed with 1.2% Avicel CL-611 in MEM containing 2% FBS, 10 mM HEPES, MEM Non-Essential Amino Acids and 100 U/ml penicillin and 100 μg/ml streptomycin. At 7 days post-infection, cells were fixed for 30 min at room temperature in 3.7% formaldehyde and permeabilized for 30 min at room temperature in 0.5% Triton X-100. Infected cells were immunostained for 2 h at room temperature with anti-NP monoclonal antibody clones TULV1 for ANDV, MAPV, TULV, PHV (a kind gift by Rainer Ulrich, Friedrich-Loeffler-Institut)_(42), B5D9 for HTNV (Progen, cat# B5D9-C), DOBV2 for THAIV (a kind gift by Rainer Ulrich, Friedrich-Loeffler-Institut) (42), A1C5 for PUUV (Progen, cat# A1C5-P). Immuno-stained foci were stained with either an IRDye 800CW-conjugated anti-mouse IgG (LI-COR, cat# 926-32210) or an HRP-conjugated anti-mouse IgG (Invitrogen, cat# 62-6520) secondary antibody. Immuno-stained foci were imaged using an Odyssey CLx imaging system (LI-COR) and analysed with the Image Studio software (LI-COR) for fluorescently labelled foci or using KPL TrueBlue Peroxidase Substrate for HRP-labelled foci. Infectious viral titres are determined as focus-forming units (FFU) per ml.

### Viral genome segment RT-PCR (vgsRT-PCR)

RNA was extracted from virus stocks, cells and tissues using TRIzol Reagent (Invitrogen) with either the TRIzol-chloroform method or the Direct-zol RNA Miniprep kit (Zymo Research) according to the manufacturers’ instructions. Reverse transcription of viral RNA was performed using the Maxima H Minus Reverse Transcriptase (Thermo Scientific) and a pair of universal hantavirus primers complementary to the conserved terminal 3’ vRNA and cRNA promoter of all hantavirus segments (Table 2). PCR amplification of entire genome segments was performed using the PrimeSTAR GXL DNA Polymerase (TaKaRa Bio) and primers complementary to the terminal 5’ and 3’ promoter regions and specific to each virus species and segment as detailed in Table 2. vgsRT-PCR reactions were resolved by 0.7% agarose gel electrophoresis and imaged on a ChemiDoc Imaging System (Bio-Rad). At Sunderby Hospital (Luleå/Sweden), reactions were resolved by 1% agarose gel electrophoresis and imaged on a E-Gel Power Snap system (Invitrogen).

**Table 2.**
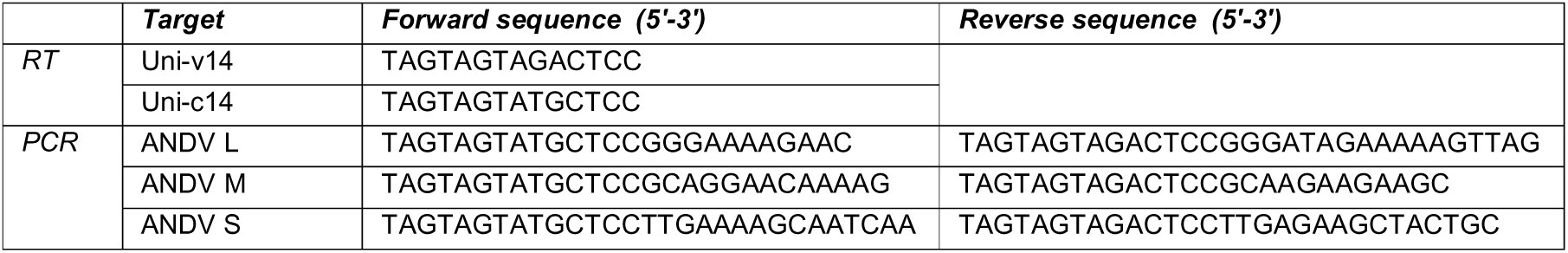
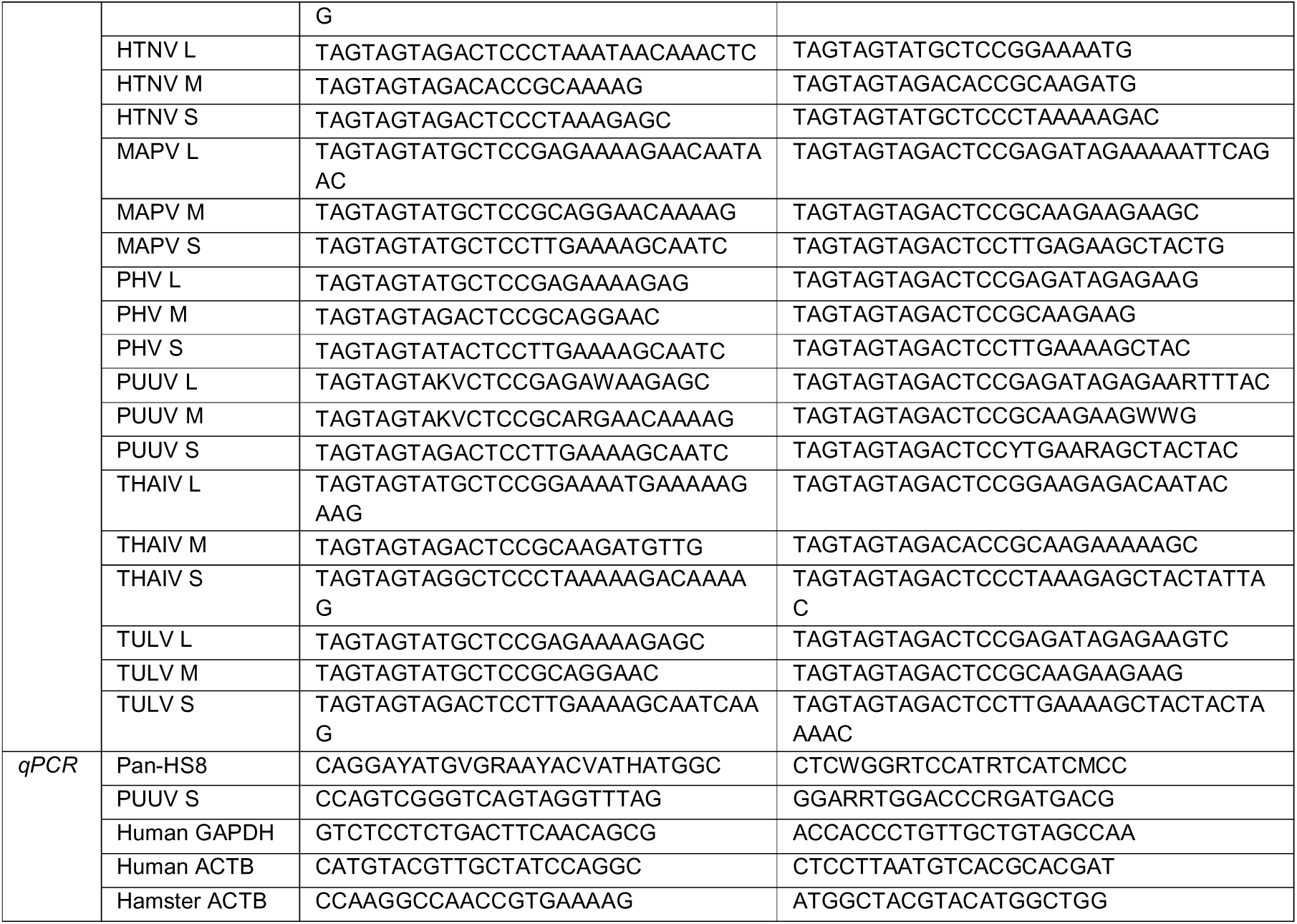
Primers used for vgsRT-PCR and RT-qPCR.

#### Quantitative real-time PCR

Cells and homogenised tissues were lysed in TRIzol Reagent (Invitrogen) and total RNA was extracted and DNase I-treated using either the Direct-zol RNA Miniprep kit (Zymo Research) or using the TRIzol-chloroform method according to the manufacturer’s instructions. Reverse transcription using oligo(dT)/random hexamer primers was performed using the PrimeScript RT Master Mix (TaKaRa Bio) according to the manufacturer’s instructions. Quantitative real-time PCR was performed using the Luna Universal qPCR Master Mix (NEB) on a LightCycler 480 II Instrument (Roche) according to the manufacturers’ instructions. Primer sequences used are detailed in Table 2. Delta-delta-cycle threshold (ΔΔCT) values were determined relative to a housekeeping gene (ACTB or GAPDH) and normalized to the mean of the indicated control group. RNA samples from infected bank voles were tested for the presence of PUUV RNA as previously described (43).

#### In vitro transcription of viral RNA

RNA was extracted from PUUV/RUS/Cg/Kazan/E6/1982 virus stocks using TRIzol Reagent (Invitrogen) and the vgsRT-PCR amplification of the S segment was performed as described above with a modified forward primer containing a T7 promoter overhang: 5’-TAATACGACTCACTATAGGGTAGTAGTAGACTCCTTGAAAAGCAATC-3’. The full-sized PCR product was gel purified using the Zymoclean Gel DNA Recovery Kit (Zymo Research) and used as a template for in vitro transcription (IVT) using the HiScribe T7 mRNA Kit with CleanCap Reagent AG (NEB). In vitro transcribed RNA was purified using the RNA Clean & Concentrator-25 kit (Zymo Research).

#### Animals and husbandry

Five 4–5-week-old female Syrian hamsters (*Mesocricetus auratus*) were purchased from Janvier labs, France, transported to the BSL-4 facility of the Robert Koch Institute and randomly assigned to three groups of five animals each. A maximum of four hamsters was housed in IVC GR900 cages (900 cm^2^, Tecniplast Sealsafe). Low-dust bedding, a shelter and nesting material were provided. Animals were acclimated for 7 days and kept in a reversed 12-h daylight cycle, at 21 °C (±1 °C) and 50% (±10%) of relative humidity. Water and food were offered *ad libitum* throughout the experiment.

One breeding pair of wild-caught bank voles (*Clethrionomys glareolus*) captured from the Sipoo region in Finland and four breeding pairs obtained from a bank vole colony at the Friedrich-Loeffler Institute in Germany (44) were used to establish a bank vole colony at the University of Helsinki. The wild-caught animals were quarantined for 3 weeks and tested for PUUV antibodies to exclude PUUV-infected individuals and all animals received antiparasitic selamectin treatment. The animals were housed in groups and provided with bedding, nesting material, play tubes and gnawing material as well as standard rodent pellets and water ad libitum.

#### Animal infections with hantaviruses

Animal experiments were performed in accordance with national regulations on animal experimentation and in conformity with the guidelines from the European Union directive 2010/63/EU.

All animal work with infectious ANDV was performed in the BSL-4 laboratory at the Robert Koch Institute. Approval for animal experiments at the Robert Koch Institute (Berlin, Germany) was granted (G 0125/23) by the Regional Office for Health and Social Affairs, Berlin. Under isoflurane anaesthesia, hamsters were weighed, ear-tagged and intranasally inoculated with 1,000 FFU (*n*=5) of ANDV Chile-9717869 in 100 µl of sterile PBS using a p200 pipette. Animals were scored and observed daily for the development of signs of disease. Animals that reached the endpoint scoring criteria were euthanized using an overdose of isoflurane and exsanguination by cardiac puncture (9-11 dpi). Sections of lung tissues were collected, weighed and homogenized in TRIzol, in a volume ratio of 1/20, using a stainless-steel bead and a TissueLyser II (QIAGEN) homogenizer and stored at −80 °C.

Experimental infections of bank voles were done according to a previously established experimental PUUV infection model (16) using adult bank voles housed in groups of 3-5 animals under BSL-3 conditions in an IsoCage biocontainment system (Scanbur). The animals were subcutaneously infected with 500 focus forming units (FFU) of PUUV strain FIN/Cg/Suonenjoki/1/2016 and euthanized by cervical dislocation at 7 days post-infection. Organs were collected into TRIzol and lung RNA was isolated according to the standard TRIzol RNA extraction protocol. All animal protocols were approved by the Animal Experimental Board of Finland, license number ESAVI/45015/2021.

RNA extracted from lung tissue of a PUUV-infected wild bank vole (*Clethrionomys glareolus*) caught in the Västerbotten region, Sweden, was kindly provided by Frauke Ecke (Swedish University of Agricultural Sciences, Sweden). Viral RNA was extracted using the NucleoSpin RNA kit (Macherey-Nagel) according to the manufacturer’s instructions.

#### Nanopore sequencing

PCR amplicons for all three segments were pooled for each sample. Pooled PCR amplicons were purified using the DNA Clean & Concentrator-25 kit (Zymo Research). Sequencing libraries were prepared using the Native Barcoding Kit 24 V14 (Oxford Nanopore Technologies; SQK-NBD114.24) and 200 fmol of PCR amplicon and sequenced on a MinION Mk1D device (Oxford Nanopore Technologies) using a MinION/GridION Flow Cell (DNA) (Oxford Nanopore Technologies; FLO-MIN114) according to the manufacturer’s instructions.

At Sunderby Hospital, Luleå no purification steps were done prior to barcoding. PCR amplicons for each of the three segments were individually barcoded using 50 ng of amplicon and then pooled for downstream processing according to the manufacturer’s instructions. Sequencing libraries with a total of 352 ng DNA were prepared using the Rapid Barcoding Kit 96 (Oxford Nanopore Technologies; SQK-RBK 110.96) and sequenced on a MinIon Mk 1B device (Oxford Nanopore Technologies) using a MinION/GridION Flow Cell (DNA) (Oxford Nanopore Technologies; FLO-MIN114) according to the manufacturer’s instructions.

#### Read processing and reference-guided genome assembly

Raw sequencing data were basecalled in super-accurate (sup) mode and demultiplexed using *Dorado* (v1.2.0) with adapter trimming enabled. Preprocessed *fastq* files were quality filtered, retaining reads with a minimum mean Phred quality score of 10, and sorted into separate size bins (S segment bin = 1600-2100 bp; M segment bin = 3400-3900 bp; L segment bin = 6200-6700 bp) with *Chopper* (v0.13.0). Selection of an appropriate hantavirus reference genome was automated by performing a mapping test of downsampled (5000 reads) raw fastq files using *Minimap2* (v2.31) (*-cx map-ont -- secondary=no*) to all generic references in Table 3 (45). The closest generic reference genome based on mapping rate and identity was then used as a reference scaffold for genome assembly. Size-filtered *fastq* read bins were separately aligned to the corresponding segment of the selected generic hantavirus reference genome using *Minimap2* (v2.31) with parameters optimized for alignment of diverse sequences (*-ax lr:hq -k 15 -w 5 -m 30 -B 3 -O 4,24 -E 2,1 --secondary=no --end-bonus=15*). SAM files were converted to BAM format, sorted, and indexed using *SAMtools* (v1.23.1) (46). BAM alignments were iteratively polished into consensus sequences and polished using *Medaka* (v2.2.2) functions *inference* and *sequence* (https://github.com/nanoporetech/medaka).

**Table 3.**
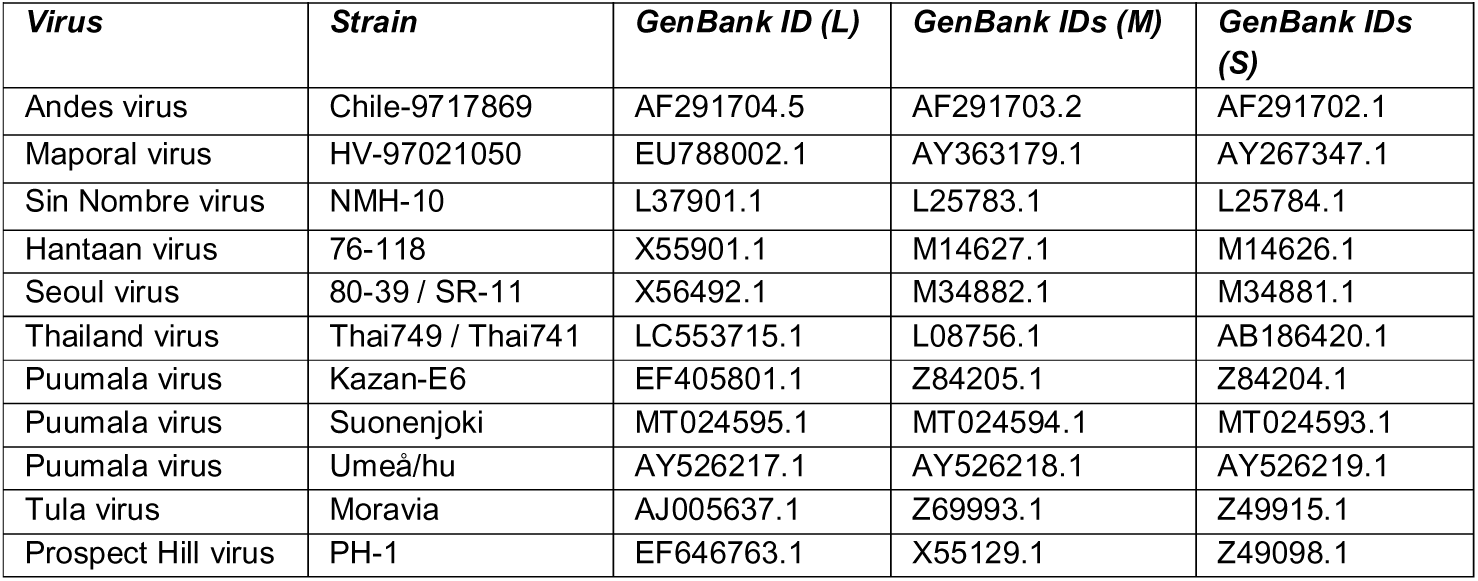
Viral reference genomes for reference selection.

| *Virus* | *Strain* | *GenBank ID (L)* | *GenBank IDs (M)* | *GenBank IDs (S)* |
| --- | --- | --- | --- | --- |
| Andes virus | Chile-9717869 | AF291704.5 | AF291703.2 | AF291702.1 |
| Maporal virus | HV-97021050 | EU788002.1 | AY363179.1 | AY267347.1 |
| Sin Nombre virus | NMH-10 | L37901.1 | L25783.1 | L25784.1 |
| Hantaan virus | 76-118 | X55901.1 | M14627.1 | M14626.1 |
| Seoul virus | 80-39 / SR-11 | X56492.1 | M34882.1 | M34881.1 |
| Thailand virus | Thai749 / Thai741 | LC553715.1 | L08756.1 | AB186420.1 |
| Puumala virus | Kazan-E6 | EF405801.1 | Z84205.1 | Z84204.1 |
| Puumala virus | Suonenjoki | MT024595.1 | MT024594.1 | MT024593.1 |
| Puumala virus | Umeå/hu | AY526217.1 | AY526218.1 | AY526219.1 |
| Tula virus | Moravia | AJ005637.1 | Z69993.1 | Z49915.1 |
| Prospect Hill virus | PH-1 | EF646763.1 | X55129.1 | Z49098.1 |

#### Genome alignment, consensus generation, and variant calling

Preprocessed *fastq* files were quality filtered with *Chopper* (v0.13.0), retaining reads with a minimum mean Phred quality score of 10 and excluding reads longer than 7,000 nucleotides to remove likely sequencing artefacts and concatemeric molecules (47). For each viral species, filtered reads were aligned against the corresponding reference genome using *Minimap2* (v2.31) with parameters optimized for DVG detection (*-ax splice:hq -C0 -- secondary=no --end-bonus=2 -un --cs --splice-flank=no -z 200,50 -Y -m 20 -s 30*), with secondary alignments being excluded while retaining supplementary alignments to facilitate downstream analyses of copy-back DVGs. SAM files were converted to BAM format, sorted, and indexed using *SAMtools* (v1.23.1) (46). Genome-wide coverage profiles were generated as BEDGRAPH files using *BEDTools* (v2.31.1), and alignments were additionally converted to BED12 format for downstream processing (48). For consensus sequence generation and variant calling, only full-length viral genome reads were used. To identify these, reference genome lengths were obtained from indexed FASTA files. Reads were retained if they consisted of a single alignment block (single exon) and aligned across at least 95% of the corresponding reference genome length. Read identifiers meeting these criteria were used to subset the sorted BAM files. Consensus genome sequences were generated and polished from the full-length read alignments using the *Medaka* (v2.2.2) functions *inference* and *sequence* (https://github.com/nanoporetech/medaka) with a minimum read depth of 5. Per-base nucleotide counts were calculated from the full-length read alignments using *bam-readcount* (v1.0.1) with a window size of 5 bases and a maximum read depth of 1,000,000 (49). Variant calling was subsequently performed using an in-house Python script (*variant_caller_v1.3*), which identified nucleotide variants from the bam-readcount output.

#### Overview of the DVG detection pipeline

Defective viral genomes (DVGs) were identified with ORCHID (ONT-based Recombination and Copy-back Haplotype Identification for DVGs), a custom Python pipeline (orchid v1.0; https://github.com/DepledgeLab/ORCHID) that takes a coordinate-sorted, indexed BAM file and the reference FASTA as input. The pipeline detects two classes of DVG. Deletion DVGs appear as large reference-skipping (N) operations within a primary alignment. Copy-back and snap-back DVGs appear as a primary alignment accompanied by a supplementary alignment on the same genome segment but on the opposite strand. Segment lengths were taken from the reference FASTA. The pipeline uses pysam, pandas, NumPy, and matplotlib. Default parameter values are listed in Table 4.

**Table 4.** Default ORCHID parameters.

| **Setting** | **Purpose** | **Default value** |
| --- | --- | --- |
| `--min-mapq` | Min mapping quality for primary and supplementary alignments | 10 |
| `--min-deletion-size` | Min N-op size (nt) or overlap size to call a deletion | 20 |
| `--min-deletion-overhang` | Min query sequence (nt) on each side of a deletion junction, including soft-clipped bases | 20 |
| `--min-deletion-support` | Min reads supporting a deletion junction | 2 |
| `--min-copy-back-overlap` | Min reference overlap (nt) between primary and supplementary alignments for a copy-back call | 20 |
| `--min-copy-back-support` | Min reads supporting a copy-back junction | 2 |

##### Read filtering and pre-classification

Only primary alignments were used to define the read population. Unmapped reads, secondary alignments, and primary alignments with a mapping quality (MAPQ) below 10 were excluded from classification, and their numbers were recorded per segment so that the output could be reconciled with the input BAM file. Supplementary alignments were not counted as separate reads but were used for copy-back detection. Each retained read was assigned, from the CIGAR string of its primary alignment, to one of three classes: (i) large-N reads, with at least one N operation of 20 nt or longer, which were candidates for deletion DVGs; (ii) small-N reads, in which N operations were present but all shorter than 20 nt, which were putative deletion DVG candidates; and (iii) no-N reads, with no N operations. Only no-N reads were eligible for copy-back detection, which ensured that no read could be counted as both a deletion and a copy-back DVG.

##### Detection of deletion DVGs

For each large-N read, the CIGAR string was traversed to obtain the reference and query coordinates of every N operation of at least 20 nt. A junction was retained if at least 20 nt of query sequence lay on each side of the N operation. These overhangs were measured in query space and therefore include soft-clipped bases. The junction start and end were the reference coordinates flanking the N operation, and the junction size was the length of the N operation.

##### Detection of copy-back DVGs

Copy-back candidates were no-N reads with a supplementary alignment (MAPQ ≥ 10) on the same segment as, and on the opposite strand to, the primary alignment; if a read had several, the one with the highest MAPQ was used. The overlap between the primary and supplementary alignments in reference coordinates was calculated. Candidates with an overlap of at least 20 nt were classified as copy-back DVGs, with the junction start, end and size defined by the boundaries and extent of the shared reference interval. Candidates with a shorter overlap, or none, were classified as putative artefacts, because opposite-strand alignments that share little or no reference sequence are inconsistent with a fold-back and may reflect PCR chimeras or spurious supplementary alignments.

##### Junction merging and support filtering

Junctions were merged across reads and strands by segment, start coordinate, end coordinate, and DVG type. For each unique junction, the pipeline reports the number of supporting reads, the mean MAPQ, and the mean arm lengths. Junctions of either type were retained only if they were supported by at least two reads. Reads whose junctions were all removed by this filter were reassigned to the corresponding putative category (putative deletion or putative copy-back), so that per-segment read totals were unchanged.

##### Read classification and DVG fraction

Every classified read was assigned to exactly one of six categories per segment: deletion, copy-back, putative deletion, putative copy-back, putative artefact, or non-DVG. The putative deletion category comprises small-N reads, large-N reads that failed the overhang filter, and reads whose junctions failed the support filter. The DVG fraction for each segment was calculated as the number of deletion and copy-back reads divided by the total number of classified reads. An internal check confirmed that the category counts summed to the total for every segment.

##### Visualization

For each segment and DVG type, junctions were plotted as arcs along a schematic of the segment (parabolic arcs for deletions, S-shaped curves for copy-backs), with arc height scaled to junction size relative to segment length. Junction start was also plotted against junction end in a scatter plot, with point size proportional to read support. Arc and point colors encode log-scaled read support. An optional track shows read depth from primary alignments. Junction size distributions were plotted as histograms with 40 equal-width bins across the segment length, and read classification was plotted as stacked bar charts of counts and proportions. The exact command line and all parameter values are written to the final page of the output PDF.

#### Multiple sequence alignment and phylogenetic tree construction

Viral sequences were either obtained from generated consensus sequences or from the National Center for Biotechnology Information (NCBI) GenBank database. GenBank accession numbers used are detailed in Table 5. Segments were aligned independently using the DECIPHER package (v3.6.0) in R, which implements the MUSCLE (Multiple Sequence Comparison by Log-Expectation) algorithm (50). All alignment parameters were set to default values. Pairwise evolutionary distances were calculated using the Jukes-Cantor (JC69) distance model implemented in the ape package in R (51). Phylogenetic trees were constructed using the neighbor-joining method as implemented in the ape package and rooted using midpoint rooting. (52).

**Table 5.**
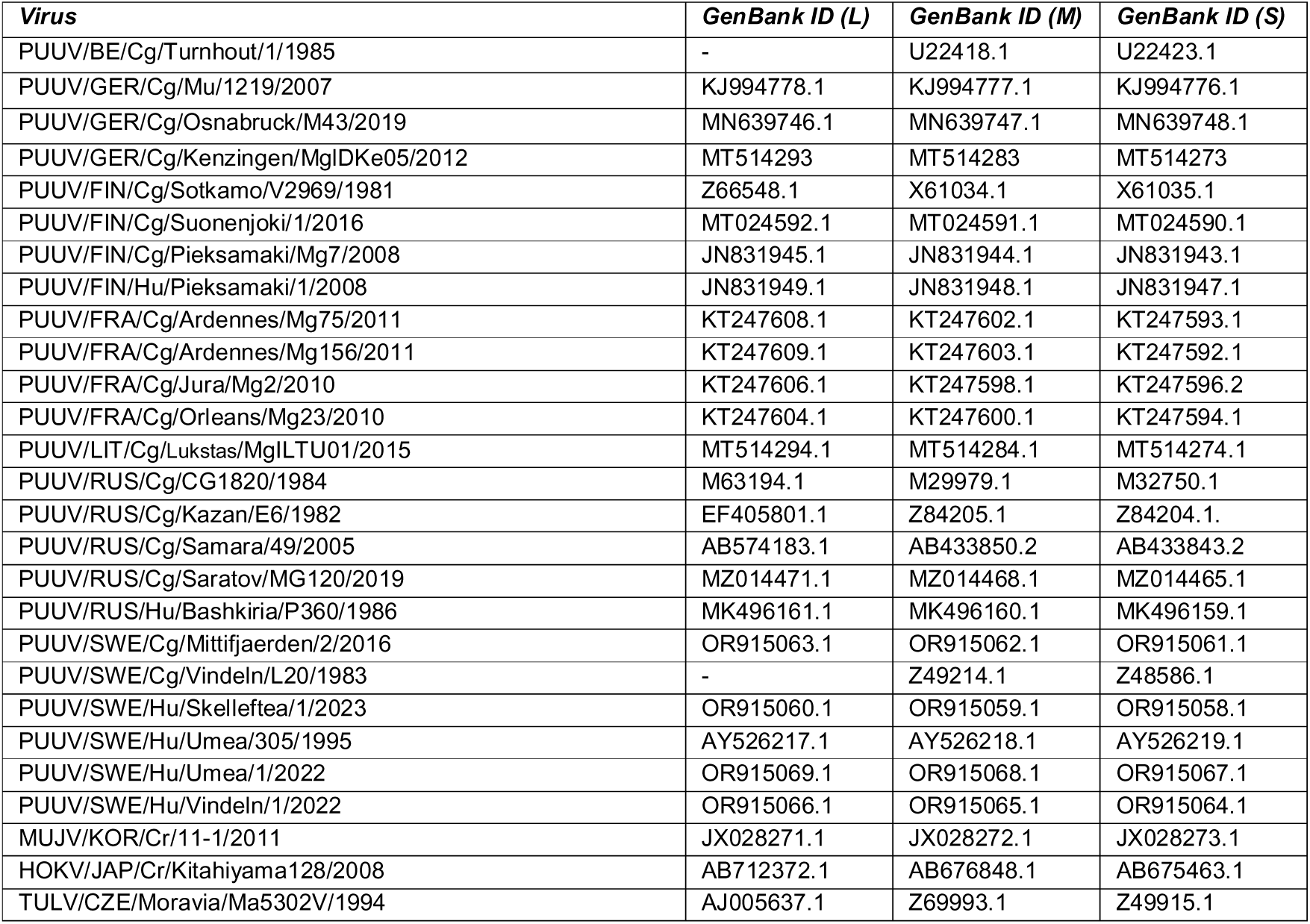
Accession numbers of sequences used for multiple sequence analysis.

## Data and code availability

The raw sequencing data for this study have been deposited in the European Nucleotide Archive (ENA) at EMBL-EBI under accession number PRJEB128616. The ORCHID source code is available at https://github.com/DepledgeLab/ORCHID.

## ACKNOWLEDGEMENTS

This study was supported by the Swedish Research Council (2023-02595 and 2024-03783 to B.E.N.), Visare Norr (RV-970009 to A.T.B), OE Edla Johansson Foundation (to A.T.B), Åke Wiberg Foundation (M24-0049 to A.T.B) and Region Västerbotten ALF (RV-1034819). D.P.D. is supported by a German Centre for Infection Research (DZIF) Associate Professorship and also receives funding from the Deutsche Forschungsgemeinschaft (DFG, German Research Foundation) under Germany’s Excellence Strategy - EXC 2155 - project number 390874280.

We acknowledge support from the European Virus Archive Global (EVA-GLOBAL) project that has received funding from the European Union’s Horizon 2020 research and innovation program under grant agreement No 871029. The bioinformatical analyses were enabled by resources provided by the National Academic Infrastructure for Supercomputing in Sweden (NAISS), partially funded by the Swedish Research Council through grant agreement no. 2022-06725. The authors also acknowledge the support of the MHH-HPC in completing this work.

